# Immune Reconstitution and Antigenic Exposure History Shape Virus-Specific Humoral Immunity in People with HIV

**DOI:** 10.64898/2026.09.26.754322

**Authors:** Juan Han, Shuyan Zhang, Chunhai Liu, Shi Zou, Lanfang Shi, Tongtong Li, Chenxia Xu, Chong Chen, Ke Liang, Qian Wang, Lihong Liu

**Affiliations:** State Key Laboratory of Virology and Biosafety, College of Life Sciences, Wuhan University, Wuhan 430072, China; TaiKang Center for Life and Medical Sciences, Wuhan University, Wuhan 430072, China; Department of Infectious Diseases, Zhongnan Hospital of Wuhan University, Wuhan 430071, China; Department of Infectious Diseases, Wuhan Jinyintan Hospital, Tongji Medical College of Huazhong University of Science and Technology, Wuhan 430023, China; Hubei Clinical Research Center for Infectious Diseases, Wuhan 430040, China; Wuhan Research Center for Communicable Disease Diagnosis and Treatment, Chinese Academy of Medical Sciences, Wuhan 430040, China; Joint Laboratory of Infectious Diseases and Health, Wuhan Institute of Virology and Wuhan Jinyintan Hospital, Chinese Academy of Sciences, Wuhan 430040, China; School of Pharmaceutical Sciences and Institute of Materia Medica, Xinjiang University, Urumqi 830017, China

**Keywords:** immune reconstitution, virus-specific humoral immunity, immune imprinting, neutralizing antibodies, antigenic evolution, people living with HIV

## Abstract

CD4^+^ T-cell recovery is widely used to assess immune reconstitution in people with HIV, but its relationship to virus-specific humoral immunity across pathogens remains unclear. We characterized neutralizing activity against SARS-CoV-2, HIV-1, and influenza viruses in ART-treated, virologically suppressed individuals spanning distinct CD4^+^ T-cell levels. BA.5 was the dominant SARS-CoV-2 neutralization target across CD4 strata, whereas D614G neutralization increased with greater immune reconstitution. This gradient was largely absent for the antigenically divergent NB.1.8.1 and XFG variants. HIV-1 neutralization was inversely associated with CD4^+^ T-cell count, while influenza showed strain-dependent patterns. Neutralization-to-protection bridging preserved a CD4-associated gradient for D614G but not for highly immune-evasive variants. These findings show that immune reconstitution does not translate uniformly into antiviral humoral function. Instead, antibody profiles reflect host immune status, prior antigenic exposure, and viral antigenic evolution, supporting pathogen-specific functional assessment alongside conventional immune monitoring.

## INTRODUCTION

Virus-specific humoral immunity is shaped by the interplay among viral antigenic properties, prior antigenic exposures and the immune memory, and the functional state of T and B cells. Human immunodeficiency virus type 1 (HIV-1) infection causes persistent perturbations of both cellular and humoral immunity. Effective antiretroviral therapy (ART) suppresses HIV-1 replication and enables progressive recovery of peripheral CD4^+^ T-cell counts in most people living with HIV (PLWH). However, the extent of immune reconstitution varies substantially among individuals, and abnormalities in B-cell activation, memory B-cell composition, and antigen-specific antibody responses can persist despite sustained virological suppression.^1^ Thus, recovery of CD4^+^ T-cell counts does not necessarily imply uniform restoration of humoral immune function. A central unresolved question is whether the degree of CD4^+^ T-cell recovery reliably predicts functional antibody immunity, or whether this relationship is conserved across pathogens with distinct antigenic and exposure histories.

SARS-CoV-2 offers a unique model for addressing this question because population immunity has been shaped within a compressed time frame by vaccination, repeated infection, and rapid antigenic evolution. In PLWH, lower current or nadir CD4^+^ T-cell counts have been associated with more severe COVID-19,^2^ and individuals with peripheral CD4^+^ T-cell counts below 200 cells/µL generally mount weaker binding and neutralizing antibody responses after SARS-CoV-2 vaccination than those with more complete immune recovery.^3,4^ However, most early studies evaluated ancestral SARS-CoV-2 or relatively early variants. Since then, the spike glycoprotein has undergone extensive antigenic evolution. Marked antibody escape emerged with Omicron BA.1,^5^ increased further in BA.4/5 and BQ/XBB lineages,^6,7^ and continued among BA.2.86/JN.1 descendants.^8,9^ More recently, NB.1.8.1 and XFG have shown enhanced escape from serum neutralizing antibodies.^10^ This progressive antigenic divergence raises a key question: does the humoral advantage associated with CD4^+^ T-cell recovery persist as circulating viruses move increasingly far from the antigens that established pre-existing immune memory?

This question is further complicated by immune imprinting. Following exposure to an antigenically related variant, pre-existing memory B cells can be preferentially recalled, biasing the antibody response toward previously encountered antigens. However, such imprinting is not immutable. Repeated Omicron exposures can alleviate ancestral-strain imprinting and generate antibodies with increased recognition of contemporary variants,^11^ while clonal analyses have shown that pre-existing memory B-cell lineages can redirect their specificity toward later Omicron variants through continued somatic evolution.^12^ Whether HIV-associated immune dysfunction alters this process remains unknown. One study reported weaker ancestral-strain imprinting and relatively stronger BA.5 neutralization in PLWH than in HIV-negative controls,^13^ whereas a more recent study of ART-treated PLWH after BA.4/5 or BF.7 breakthrough infection found persistent preferential neutralization of ancestral D614G.^14^ These apparently divergent observations suggest that immune imprinting in PLWH may depend on the degree of immune reconstitution, vaccine platform, infecting lineage, and cumulative exposure history, rather than on HIV status alone.

More broadly, the relationship between current CD4^+^ T-cell status and humoral immunity may differ fundamentally across viruses. HIV-1 provides persistent antigenic stimulation during chronic infection and can drive prolonged virus-antibody coevolution, including the gradual development of neutralization breadth in a subset of individuals.^15^ Influenza viruses, by contrast, repeatedly reshape antibody repertoires through seasonal infection, vaccination, antigenic drift, and recall of pre-existing immunity; antibody-landscape studies have demonstrated how these cumulative exposures generate strain-dependent serological profiles across decades of viral evolution.^16^ SARS-CoV-2 represents yet another regime, characterized by rapid sequential vaccination and infection superimposed on unusually fast antigenic change. These distinct evolutionary and exposure trajectories predict that the same host immune state may map onto different functional antibody phenotypes for different pathogens. Peripheral CD4^+^ T-cell counts may therefore be informative for some antigenic contexts but insufficient as a universal surrogate of pathogen-specific humoral competence.

Here, we examined ART-treated, virologically suppressed PLWH with markedly different levels of CD4^+^ T-cell recovery to determine how host immune reconstitution interacts with antigenic evolution and exposure history to shape virus-specific humoral immunity. We leveraged the relatively structured SARS-CoV-2 exposure history in China, where ancestral-strain inactivated vaccination was followed by widespread BA.5.2/BF.7 and related Omicron infections and subsequent exposure to later Omicron descendants. We first profiled serum neutralization against 22 SARS-CoV-2 variants and four related sarbecoviruses in PLWH with CD4^+^ T-cell counts below 200 or above 500 cells/µL, integrating spike-binding measurements with antigenic cartography to resolve immune-imprinting patterns (**Figure 1**). We then validated the major SARS-CoV-2 neutralization phenotypes in an expanded cohort of 264 PLWH and 91 HIV-negative controls (**Figure 2**), and compared HIV-1 and influenza virus neutralization in the same population to test whether associations with CD4^+^ T-cell status were conserved across pathogens (**Figure 3**). Finally, because neutralizing antibody levels can serve as a quantitative correlate of protection against SARS-CoV-2,^17^ we used neutralization-protection immune bridging to estimate the protective implications of the observed antibody profiles (**Figure 4**). We found that BA.5-directed neutralization remained relatively preserved across levels of immune reconstitution, whereas neutralization of ancestral D614G increased with higher CD4^+^ T-cell counts; notably, this association was markedly attenuated for the antigenically advanced NB.1.8.1 and XFG variants. In contrast, HIV-1 neutralization was generally inversely associated with CD4^+^ T-cell counts, while influenza neutralization showed pronounced strain dependence. Together, these findings establish that current CD4^+^ T-cell status does not map uniformly onto functional humoral immunity. Instead, its relationship with virus-specific antibody activity is jointly conditioned by prior antigenic exposure, immune memory, and the evolutionary trajectory of the virus.

**Figure 1.**
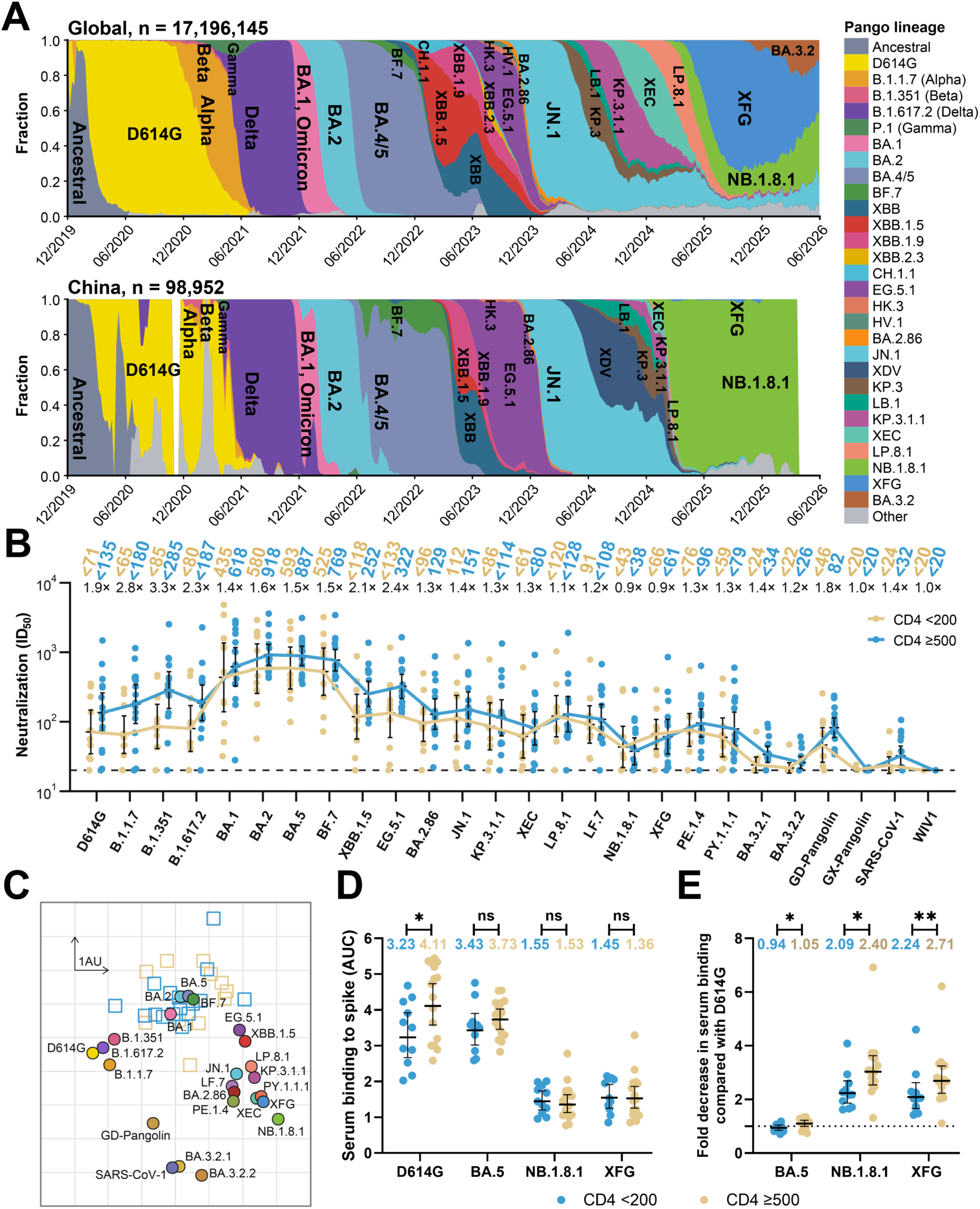
Neutralization and binding across SARS-CoV-2 variants in people living with HIV (PLWH). **A.** Frequencies of SARS-CoV-2 Pango lineages globally and in China from December 2019 to June 2026. Sequences were obtained from the Global Initiative on Sharing All Influenza Data (GISAID). *n* indicates the total number of sequences in each dataset. **B.** Serum neutralization against 22 SARS-CoV-2 variants and four related sarbecoviruses in PLWH with peripheral CD4^+^ T-cell count <200 cells/μL (CD4<200; *n* = 11) or ≥500 cells/μL (CD4≥500; *n* = 16). Lines connect geometric mean ID_50_ titers (GMTs). GMTs for each group and fold differences between groups (CD4 ≥500/CD4 <200) are shown above the plot; GMT are labeled with “<” when at least one sample within the group was at or below the limit of detection. The dashed horizontal line denotes the assay detection limit (ID_50_ = 20). **C.** Antigenic map generated from the neutralization data in panel **B**. Squares denote serum samples, colored by CD4^+^ T-cell group, and circles denote viruses. One grid interval represents one antigenic unit (AU), corresponding to an approximately twofold difference in ID_50_ titer. **D.** Serum binding to spike proteins from D614G, BA.5, NB.1.8.1, and XFG, quantified as the area under the ELISA binding curve (AUC). **E.** Fold change in serum binding to BA.5, NB.1.8.1, and XFG spike proteins relative to D614G. Comparisons between the CD4 <200 and CD4 ≥500 groups were performed using unpaired t tests. ns, not significant; \**p* < 0.05; \*\**p* < 0.01. In panel **B**, **D**, and **E**, points represent individual serum samples. Geometric means are shown above the plots. Horizontal lines and error bars indicate geometric means and 95% CIs, respectively. See also Figures S1–S3 and Table S1.

**Figure 2.**
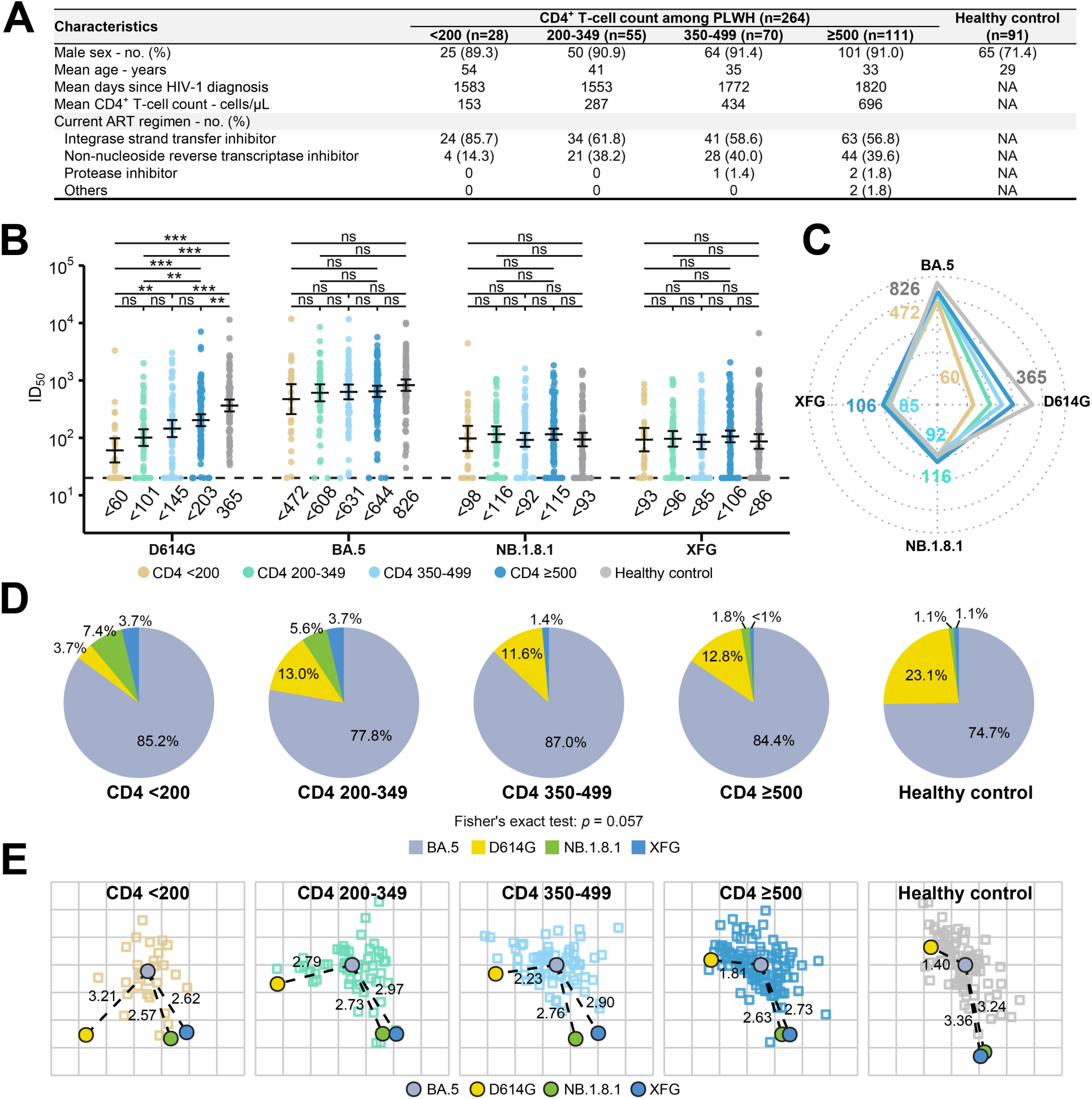
CD4^+^ T cell count-dependent neutralization of SARS-CoV-2 variant in PLWH. **A.** Clinical characteristics of five cohorts comprising 264 PLWH and 91 healthy controls. PLWH were stratified according to peripheral CD4^+^ T-cell count at sampling: <200 cells/μL (*n* = 28), 200–349 cells/μL (*n* = 55), 350–499 cells/μL (*n* = 70), and ≥500 cells/μL (*n* = 111). ART, antiretroviral therapy. **B.** Serum neutralization against D614G, BA.5, NB.1.8.1, and XFG. Points represent individual serum samples, and horizontal lines with error bars indicate GMTs and 95% CIs. The dashed horizontal line denotes the assay detection limit (ID_50_ = 20), and GMTs are shown below the plot; GMT are labeled with “<” when at least one serum within the group was at or below the detection limit. Pairwise comparisons were performed using two-sided Wilcoxon rank-sum tests with Benjamini–Hochberg adjustment. ns, not significant; \*\**p* < 0.01; \*\*\**p* < 0.001. **C.** Radar plots showing the neutralization GMT profiles of the five cohorts against D614G, BA.5, NB.1.8.1, and XFG. Line colors match the cohort colors used in panel **B**, and numbers beside each axis indicate the minimum and maximum group-level GMTs for each variant across the five cohorts. The radial axis shows ID_50_ on a log_10_ scale from 20 to 1,000. **D.** Proportions of serum samples in each cohort whose highest ID_50_ titer was directed against D614G, BA.5, NB.1.8.1, or XFG (dominant variant). Sera with all four ID_50_ values at or below the detection limit (*n* = 5) were excluded. Differences in the distributions across cohorts were assessed using Fisher’s exact test with Monte Carlo simulation (*p* = 0.057). **E.** Antigenic maps constructed from the neutralization data in panel **B**. Squares denote serum samples and circles denote variants. Numbers indicate antigenic distances from BA.5 to the other variants; one antigenic unit corresponds to an approximately twofold difference in ID_50_ titer.

**Figure 3.**
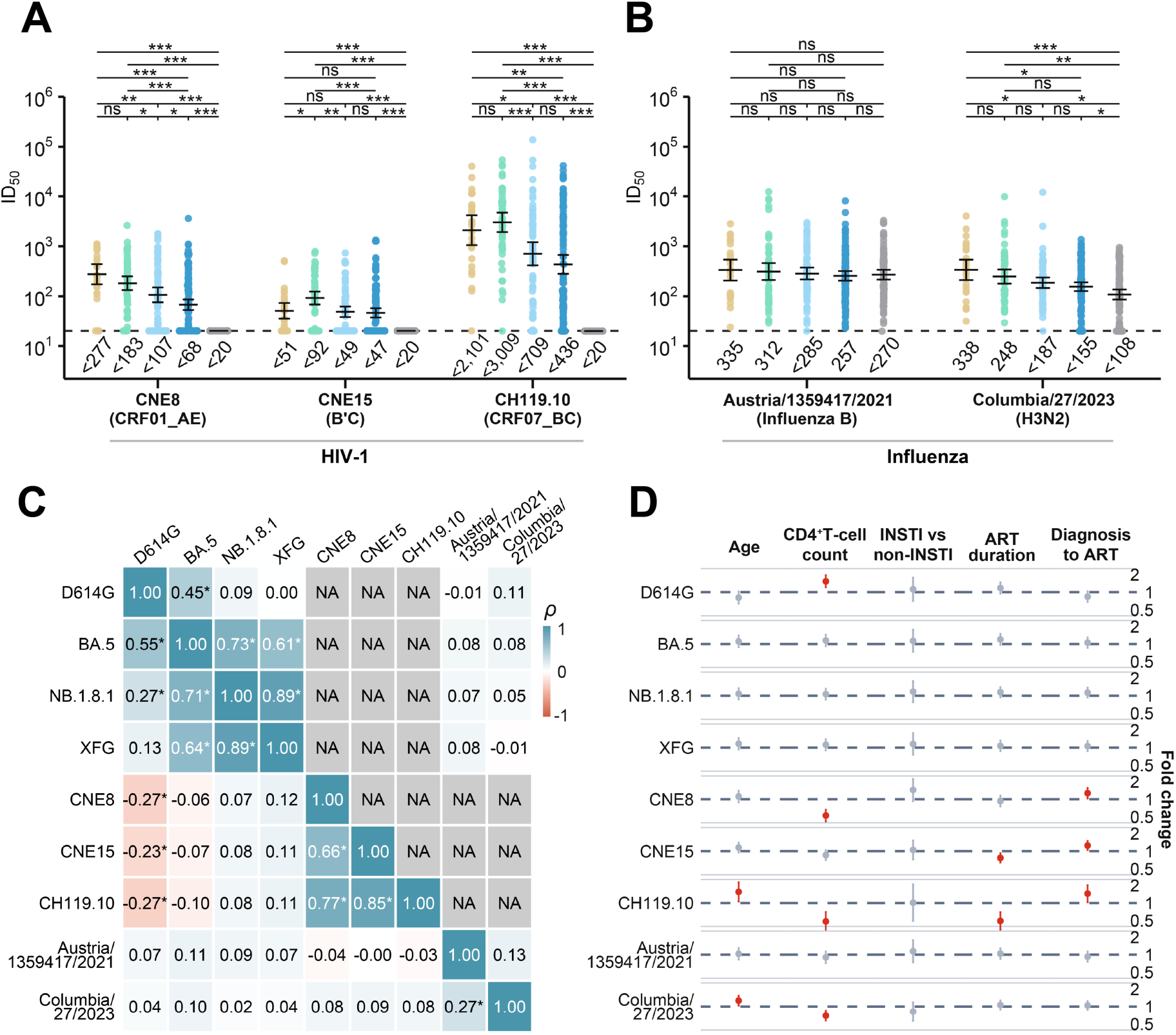
Virus-specific neutralization profiles and clinical correlates in PLWH. **A.** Serum neutralization against HIV-1 pseudoviruses CNE8 (CRF01_AE), CNE15 (BC), and CH119.10 (CRF07_BC). **B.** Serum neutralization against influenza pseudoviruses B/Austria/1359417/2021 and A/Columbia/27/2023 (H3N2). **C.** Spearman correlation matrices of log_10_-transformed ID_50_ values for the nine viruses; the lower triangle shows PLWH (*n* = 264) and the upper triangle shows healthy controls (*n* = 91). Cell color and the displayed value indicate Spearman’s *ρ*. NA indicates correlations that could not be estimated because healthy-control titers against HIV-1 were at or below the assay detection limit. Asterisks indicate Benjamini–Hochberg-adjusted *p* < 0.05. **D.** Multivariable linear regression analyses of associations between clinical variables and log_10_-transformed ID_50_ values in PLWH with complete clinical data (*n* = 262). Points indicate adjusted fold changes in ID_50_ and horizontal bars indicate 95% CIs. The dashed vertical line denotes a fold change of 1. Estimates shown in red have *p* < 0.05. In panels **A** and **B**, points represent individual serum samples, and horizontal lines with error bars indicate GMTs and 95% CIs. The dashed horizontal line denotes the assay detection limit (ID_50_ = 20), and GMTs are shown below the plots; GMT are labeled with “<” when at least one sample within the group was at or below the detection limit. Pairwise comparisons were performed using two-sided Wilcoxon rank-sum tests with Benjamini–Hochberg adjustment. ns, not significant; \**p* < 0.05; \*\**p* < 0.01; \*\*\**p* < 0.001. See also Figures S1, S4, and S5.

**Figure 4.**
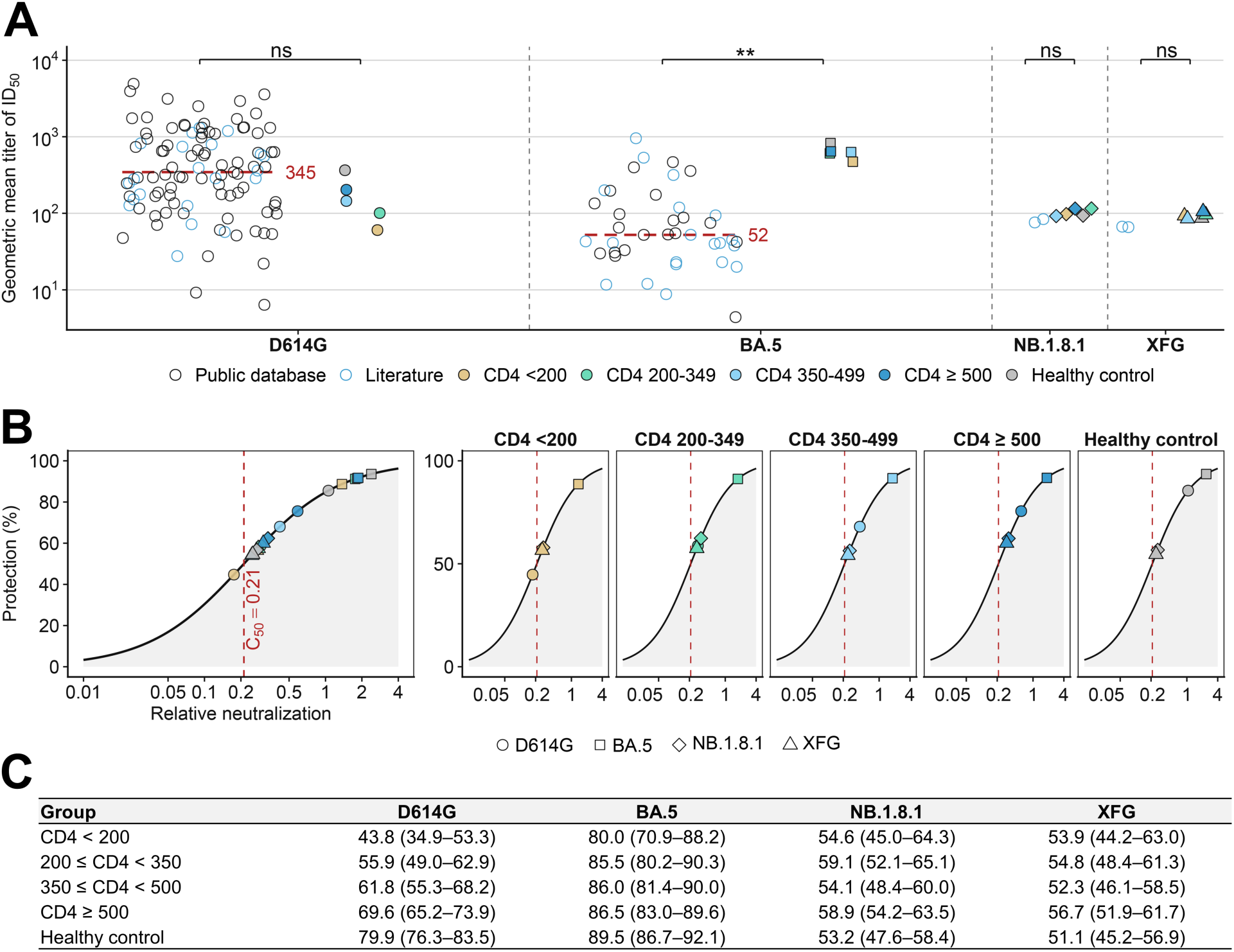
Model-based protection estimates derived from SARS-CoV-2 neutralization bridging. **A.** Curated public neutralization records from Chinese serum or plasma samples (111 cohort-level entries from 67 published studies: 83 entries covering 57 studies from the Stanford COVID-DRDB database and 28 entries extracted from 11 additional publications) together with GMTs from the five cohorts of this study for D614G, BA.5, NB.1.8.1, and XFG. Open circles denote public GMT records (black, database-derived; blue, literature-derived), and filled symbols denote cohort GMTs. Red dashed lines indicate the public median GMT of D614G (345; 103 entries) and BA.5 (52; 40 entries). Statistical analyses were performed using two-sided Mann– Whitney U tests between the four PLWH groups and all public cohorts (healthy controls not included): D614G, *p* = 0.058; BA.5, *p* = 0.0024; NB.1.8.1 and XFG, *p* = 0.133 (public reference limited to a single study). **B.** Model-based protection against symptomatic SARS-CoV-2 infection estimated by fitting relative neutralization to the logistic neutralization–protection model of Khoury et al.^17^ The left panel shows the complete relationship together with all cohort estimates; the five panels on the right show variant-specific estimates for D614G, BA.5, NB.1.8.1, and XFG across the PLWH and healthy-control cohorts. The red dashed line indicates the relative neutralization level at which the model predicts 50% protection, termed C_50_ (= 0.21). These values are model-based protection estimates rather than directly observed vaccine effectiveness or clinical efficacy. Relative neutralization is the cohort GMT divided by the public D614G GMT anchor (345). **C.** Mean model-based protection estimates for the five cohorts in this study against the indicated variants (95% CIs in parentheses). Neutralization values were normalized to the public D614G GMT anchor (345), and protection was estimated using 2,000 bootstrap resamples. **See also Table S2.**

## RESULTS

### CD4^+^ T-cell status is associated with the potency and breadth of SARS-CoV-2 neutralizing antibody responses in PLWH

To investigate how immune status influences SARS-CoV-2 humoral immunity in PLWH, we collected serum samples between May and July 2026 from 27 ART-treated PLWH with suppressed plasma HIV-1 RNA levels (≤200 copies/mL; **Table S1**). Participants were stratified according to peripheral CD4^+^ T-cell count into a severely immunocompromised group with <200 cells/μL (CD4<200, *n* = 11) and a group with relatively preserved immune reconstitution with ≥500 cells/μL (CD4≥500, *n* = 16). To capture the antigenic spectrum of SARS-CoV-2 encountered over the course of the pandemic, we first analyzed the temporal distribution of major Pango lineages globally and in China from December 2019 through June 2026 (**Figure 1A**)^18,19^. On the basis of these epidemiological trajectories, we selected SARS-CoV-2 variants spanning D614G, Alpha (B.1.1.7), Beta (B.1.351), Delta (B.1.617.2), and 18 Omicron-lineage viruses. Four related sarbecoviruses were additionally included to assess cross-sarbecovirus neutralization: the SARS-CoV-2-like GD-Pangolin and GX-Pangolin viruses, and the more divergent SARS-CoV-1 and SARS-CoV-1-like WIV1 viruses. Single-round VSV pseudoviruses bearing the corresponding spike proteins were generated for neutralization assays (**Figure S1A**)^20,21^.

Because all PLWH were receiving ART at the time of sampling, we first excluded the possibility that residual antiretroviral drugs in serum nonspecifically inhibited VSV pseudovirus infection. VSV pseudoviruses bearing the Gn/Gc glycoproteins of severe fever with thrombocytopenia syndrome virus (SFTSV; G/K/2012) or HIV-1 CH119.10 envelope were therefore used as controls. To improve HIV-1 envelope incorporation and pseudovirus infectivity, the transmembrane and cytoplasmic domains of CH119.10 Env were replaced with the corresponding regions of VSV-G, generating a chimeric envelope. This modification increased pseudovirus infectivity by approximately 82-fold relative to the wild-type construct (**Figures S1B and S1C**). Sera from nine individuals with prior SFTSV infection potently neutralized VSV-G/K/2012 pseudovirus with geometric mean neutralizing titer (GMT) of 12,471 but showed no detectable neutralization of the HIV-1 pseudovirus (GMT <20). Conversely, sera from PLWH with different ART regimens showed no detectable neutralization of VSV-G/K/2012, while retaining substantial neutralizing activity against the HIV-1 pseudovirus (**Figure S2**). These controls indicate that the neutralizing activity measured in PLWH sera was virus specific and was not attributable to nonspecific inhibition by residual antiviral drugs.

We next profiled serum neutralization across the 26-coronavirus panel (**Figure 1B**). A clear hierarchy of neutralization sensitivity emerged. Among the SARS-CoV-2 variants tested, early Omicron-lineage viruses, particularly BA.1, BA.2, BA.5, and BF.7, were neutralized most efficiently. By comparison, neutralization of D614G, B.1.1.7, B.1.351, and B.1.617.2 remained readily detectable but was lower than that of the early Omicron variants. Neutralizing activity subsequently declined across later Omicron descendants, with the contemporary NB.1.8.1 and BA.3.2-lineage variants showing among the strongest escape phenotypes in the panel. Cross-neutralization of more divergent sarbecoviruses was limited. GD-Pangolin remained partially susceptible, with GMTs of approximately <46 and 82 in the CD4 <200 and CD4 ≥500 groups, respectively. SARS-CoV-1 was only weakly neutralized, whereas GX-Pangolin and WIV1 were largely resistant. Thus, the serum antibody repertoire retained some cross-sarbecovirus activity, but this breadth was substantially weaker than neutralization of SARS-CoV-2 variants.

CD4^+^ T-cell status was associated with the overall potency of SARS-CoV-2 neutralization. For 20 of the 22 SARS-CoV-2 variants, sera from the CD4 ≥500 group showed GMTs approximately 1.1- to 3.3-fold higher than those from the CD4 <200 group. The two exceptions were XFG and NB.1.8.1, against which neutralizing titers were uniformly low and comparable between groups. The CD4 ≥500 group also showed stronger neutralization of GD-Pangolin and SARS-CoV-1. These data indicate that preserved CD4^+^ T-cell immunity is associated with greater neutralizing antibody potency and breadth, although this advantage becomes less evident against highly immune-evasive contemporary variants.

Antigenic cartography further resolved these relationships (**Figure 1C**). Early Omicron variants, including BA.1, BA.2, BA.5, and BF.7, clustered relatively close to the serum samples, whereas D614G, B.1.1.7, B.1.351, and B.1.617.2 occupied somewhat more distant positions. Variants emerging after the BF.7 period showed progressively greater displacement. NB.1.8.1 and the BA.3.2-lineage viruses were among the most antigenically distinct, lying approximately 4.5–5.3 antigenic units (AU) from the serum cluster, comparable to the distances observed for GD-Pangolin (3.8 AU) and SARS-CoV-1 (5.0 AU). Thus, the antigenic map recapitulated the neutralization hierarchy and illustrated the progressive divergence of recent SARS-CoV-2 variants from the antibody repertoire represented in these sera.

We next asked whether the preferential neutralization of early Omicron variants was also reflected in spike-binding antibody responses. Serum binding to purified spike ectodomains from D614G, BA.5, NB.1.8.1, and XFG was quantified by ELISA (**Figures 1D and S3**). In the CD4 ≥500 group, binding to D614G was significantly higher than that in the CD4 <200 group, whereas the difference for BA.5 was smaller and not statistically significant. Binding to XFG and NB.1.8.1 was low and comparable between the two groups. Within the CD4 ≥500 group, D614G elicited the strongest binding, followed by BA.5, whereas XFG and NB.1.8.1 were recognized less efficiently. In contrast, sera from the CD4 <200 group showed their strongest binding to BA.5, followed by D614G. Binding to BA.5 was largely preserved relative to D614G, whereas binding to XFG and NB.1.8.1 was reduced (**Figure 1E**). The reductions against XFG and NB.1.8.1 were approximately 2.09- to 2.71-fold in AUC, with greater losses generally observed in the CD4 ≥500 group. Together, the neutralization and binding data reveal distinct serological footprints according to CD4^+^ T-cell status: individuals with preserved CD4^+^ T-cell count retained stronger binding of the ancestral spike, whereas those with severe CD4^+^ T-cell depletion showed a relatively greater bias toward BA.5. At the neutralization level, however, both groups displayed their highest activity against early Omicron viruses.

### Expanded cohort analysis reveals CD4-dependent neutralization of earlier SARS-CoV-2 variants but convergence against contemporary variants

To validate the association between CD4^+^ T-cell status and SARS-CoV-2 neutralization in a larger population, we expanded the analysis to 264 ART-treated PLWH with plasma HIV-1 RNA ≤200 copies/mL and 91 healthy controls. PLWH were stratified into four groups according to peripheral CD4^+^ T-cell count: <200 (*n* = 28), 200–349 (*n* = 55), 350–499 (*n* = 70), and ≥500 cells/μL (*n* = 111) (**Figure 2A**). Neutralizing activity was evaluated against D614G, BA.5, NB.1.8.1, and XFG. Across all five cohorts, BA.5 was the most efficiently neutralized virus (**Figures 2B and 2C**). GMTs of BA.5 increased from 472 in the CD4 <200 group to 608, 631, and 644 in the 200–349, 350–499, and ≥500 groups, respectively, and reached 826 in healthy controls. Consistent with the GMT analysis, BA.5 represented the virus with the highest individual ID_50_ in more than 74% of samples in each cohort (**Figure 2D**). Although BA.5 neutralization showed a numerical increase with CD4^+^ T-cell count, this relationship was relatively modest.

A stronger CD4^+^ T-cell count-dependent pattern was observed for D614G. Neutralizing activity against D614G increased progressively across the CD4 strata, with the CD4 ≥500 group and healthy controls showing approximately 3.4-fold and 6-fold higher activity, respectively, than the CD4 <200 group. Notably, the relative position of D614G within the neutralization hierarchy also differed according to immune status. In individuals with CD4^+^ T-cell count ≥350 cells/μL and in healthy controls, D614G was neutralized more efficiently than NB.1.8.1 and XFG. In contrast, in the two lower-CD4^+^ T-cell count groups, neutralization of D614G fell to levels comparable to or below those of the two contemporary variants. The CD4-dependent gradient, evident for D614G and to a lesser extent BA.5, was largely absent for NB.1.8.1 and XFG. Neutralizing titers against both variants remained low across all PLWH and healthy controls (**Figures 2B and 2C**). Thus, the enhanced neutralizing capacity associated with higher CD4^+^ T-cell count was most apparent against antigenically earlier SARS-CoV-2 viruses but was substantially diminished against highly evolved contemporary variants.

Antigenic maps generated separately for the five cohorts supported this interpretation (Figure 2E). BA.5 remained the antigenically closest of the four tested viruses across groups. The antigenic distance between D614G and BA.5 decreased progressively with increasing CD4^+^ T-cell count, from approximately 3.21 AU in the CD4 <200 group to 1.40 AU in healthy controls. In contrast, the distances of NB.1.8.1 and XFG from BA.5 were comparatively stable among PLWH, generally ranging from approximately 2.57 to 2.97 AU. In healthy controls, these distances were larger, reaching approximately 3.24 AU for NB.1.8.1 and 3.36 AU for XFG.

Taken together, these data reveal an exposure-dependent neutralization pattern related to immune imprinting.^22–24^ Individuals with better-preserved CD4^+^ T-cell immunity maintained stronger responses and deeper immune memory to previously encountered variants, particularly D614G and BA.5, whereas neutralization of the more antigenically divergent XFG and NB.1.8.1 remained uniformly limited. Conversely, individuals with lower CD4^+^ T-cell count showed weaker responses to earlier variants and correspondingly less separation between ancestral and contemporary variants.

### Neutralizing antibody responses to different viruses show distinct relationships with CD4^+^ T-cell status

We next asked whether the association between CD4^+^ T-cell status and SARS-CoV-2 neutralization extended to other viral infections. We therefore measured neutralizing activity against three HIV-1 strains and two influenza viruses in the same expanded cohorts. The HIV-1 panel comprised CNE8 (CRF01_AE), CNE15 (BC)), and CH119.10 (CRF07_BC), representing lineages that have circulated globally or in China (**Figure S4A**)^25,26^. Chimeric VSV pseudoviruses were generated using the same strategy described above, with the transmembrane and cytoplasmic regions of the HIV-1 envelopes replaced by the corresponding VSV-G regions to improve infectivity (**Figures S1B and S1C**). For influenza, B/Austria/1359417/2021 and A/Columbia/27/2023 (H3N2) were selected to represent recently circulating influenza B and H3N2 lineages (**Figure S4B**)^27,28^, and VSV pseudoviruses were generated using unmodified HA and NA proteins (**Figure S1D**).

As expected, sera from healthy controls showed no detectable neutralization of the three HIV-1 pseudoviruses (**Figure 3A**). Among PLWH, CH119.10 was the most efficiently neutralized HIV-1 strain, with GMTs across the four CD4 groups ranging from approximately 436 to 3,009. CNE8 showed intermediate sensitivity, with GMTs of approximately 68–277, whereas CNE15 was the least sensitive, with GMTs of approximately 47–92. Unexpectedly, the relationship between CD4^+^ T-cell count and HIV-1 neutralization was opposite to that observed for ancestral SARS-CoV-2. Neutralizing activity against the HIV-1 pseudoviruses generally increased as CD4^+^ T-cell count decreased. However, the CD4 <200 group did not consistently show the highest titers; for CNE15 and CH119.10, its GMTs fell between those observed in the 200–349 and 350–499 groups. Thus, HIV-1-specific neutralization exhibited an inverse rather than monotonic positive relationship with contemporaneous CD4^+^ T-cell count. A virus-specific pattern was also observed for influenza (**Figure 3B**). Neutralization of influenza B/Austria/1359417/2021 did not differ substantially across the CD4^+^ T-cell strata, whereas neutralization of A/Columbia/27/2023 (H3N2) tended to increase as CD4^+^ T-cell count decreased, resembling the pattern observed for HIV-1 rather than that for D614G.

To determine whether neutralizing responses to different viruses were coordinated, we performed pairwise Spearman correlation analyses across the nine viruses (**Figure 3C**). In PLWH, neutralizing activities were most strongly correlated within viral families. Among SARS-CoV-2 variants, NB.1.8.1 and XFG showed the strongest correlation (Spearman’s *ρ* = 0.89), while BA.5 correlated with both NB.1.8.1 (*ρ* = 0.71) and XFG (*ρ* = 0.64). Neutralizing responses to CNE8, CNE15, and CH119.10 were also strongly correlated with one another (*ρ* = 0.66–0.85), whereas responses to the two influenza viruses showed a weaker but significant positive correlation (*ρ* = 0.27). In contrast, correlations across different viral families were weak or absent. One notable exception was D614G, whose neutralization showed weak inverse correlations with the three HIV-1 strains (*ρ* approximately −0.23 to −0.27). In healthy controls, the four SARS-CoV-2 variants showed a correlation pattern similar to that observed in PLWH. Correlations involving HIV-1 could not be reliably estimated in healthy controls because neutralizing titers were undetectable.

We next assessed whether the observed neutralization patterns remained associated with clinical variables after multivariable adjustment (**Figure 3D**). Data were analyzed on 262 PLWH whose complete clinical data were available. After log transformation and standardization, each one-standard-deviation increase in CD4^+^ T-cell count was independently associated with a 1.38-fold increase in D614G ID_50_ (95% CI, 1.15–1.65; *p = 4.8 × 10*^−4^). In contrast, higher CD4^+^ T-cell count was associated with lower neutralization of CNE8, CH119.10, and H3N2, with adjusted ID_50_ fold changes of 0.62, 0.58, and 0.76, respectively. The interval between HIV-1 diagnosis and ART initiation was positively associated with neutralization of all three HIV-1 strains: each one-standard-deviation increase in this interval was associated with 1.20-, 1.18-, and 1.32-fold higher ID_50_ values against CNE8, CNE15, and CH119.10, respectively. By contrast, longer ART duration was associated with reduced neutralization of CNE15 and CH119.10, with adjusted fold changes of 0.82 and 0.59. Older age was independently associated with higher neutralization of CH119.10 and H3N2. No significant associations were detected between INSTI-based versus non-INSTI-based ART regimens and neutralization of any of the nine viruses. Likewise, BA.5, NB.1.8.1, XFG, and influenza B neutralization showed no independent association with the clinical variables examined. The same directions were evident in univariate Spearman correlation analyses (**Figure S5**): CD4^+^ T-cell count correlated positively with D614G neutralization (*ρ* = 0.30) and negatively with neutralization of all three HIV-1 strains (*ρ* = −0.39 to −0.22), and age correlated positively with HIV-1 and H3N2 neutralization.

Collectively, these analyses demonstrate that immune impairment does not uniformly reduce virus-specific humoral immunity. Instead, the relationship between CD4^+^ T-cell status and neutralizing antibody activity differs markedly across viruses, consistent with contributions from exposure history, chronic infection, immune reconstitution, and virus-specific antigenic properties.

### Neutralization bridging predicts CD4^+^ T-cell count-dependent protection against historical but not contemporary SARS-CoV-2 variants

Finally, we examined how the observed differences in SARS-CoV-2 neutralization might translate into model-based protection from symptomatic infection. We curated 111 cohort-level neutralization entries from 67 published studies of Chinese serum or plasma samples (83 entries covering 57 studies from the Stanford COVID-DRDB database and 28 entries extracted from 11 additional publications) and compared these external data with GMTs from the cohorts of this study (**Table S2 and Figure 4A**). Cohort GMTs of D614G did not differ significantly from the public distribution (two-sided Mann–Whitney U test, *p* = 0.058), whereas cohort GMTs of BA.5 were significantly higher than the public reference (*p* = 0.0024); for NB.1.8.1 and XFG, the public reference comprised only two entries from a single study,^29^ precluding a meaningful comparison (*p* = 0.133). The median of the 103 available public D614G GMT values (345.4) was used as the external neutralization anchor. The D614G GMT in the healthy-control cohort was close to this reference value, supporting its use as a calibration background for neutralization bridging. Relative neutralization levels for D614G, BA.5, NB.1.8.1, and XFG were then mapped onto the logistic neutralization–protection model of Khoury et al.^17^ to estimate protection against symptomatic SARS-CoV-2 infection (**Figures 4B and 4C**). For D614G, estimated protection increased progressively with CD4^+^ T-cell count. Mean model-estimated protection was 43.8% (95% CI, 34.9–53.3%) in the CD4 <200 group, 55.9% (49.0–62.9%) in the 200–349 group, 61.8% (55.3–68.2%) in the 350–499 group, and 69.6% (65.2–73.9%) in the CD4 ≥500 group, reaching 79.9% (76.3–83.5%) in healthy controls. BA.5 yielded the highest protection estimates across all cohorts, ranging from 80.0% to 89.5%, consistent with the uniformly high neutralizing titers against this variant. In contrast, estimates for NB.1.8.1 remained within a relatively narrow range of 53.2– 59.1%, and those for XFG ranged from 51.1% to 56.7%. Unlike D614G, neither NB.1.8.1 nor XFG showed a monotonic increase in estimated protection across increasing CD4^+^ T-cell strata.

These analyses therefore identify two distinct patterns. Protection estimates against D614G retained a clear CD4-associated gradient, while BA.5 remained uniformly high because of strong neutralization across cohorts. By contrast, the extensive antigenic escape of NB.1.8.1 and XFG compressed differences between CD4 groups and produced lower protection estimates. Importantly, these values represent model-based estimates derived from relative neutralization and an external neutralization–protection relationship; they should not be interpreted as directly observed infection rates, vaccine effectiveness, or individual-level protection.

## DISCUSSION

This study examined how CD4^+^ T-cell reconstitution relates to humoral immunity across pathogens and antigenic settings in virologically suppressed PLWH. Higher CD4^+^ T-cell count was associated with stronger neutralization of most historical SARS-CoV-2 variants, whereas this gradient diminished with increasing viral divergence and was minimal for NB.1.8.1 and XFG (**Figures 1 and 2**). BA.5 was consistently the dominant neutralization target across CD4 strata, while D614G showed a stronger dependence on the degree of host recovery (**Figures 1 and 2**). By contrast, HIV-1 neutralizing activity generally increased as CD4^+^ T-cell count decreased, H3N2 showed a similar tendency, and influenza B exhibited little dependence on CD4 status (**Figure 3**). Neutralization-to-protection bridging likewise preserved a clear CD4-associated gradient for D614G but not for highly divergent contemporary SARS-CoV-2 variants (**Figure 4**). These observations do not support a model in which the extent of immune reconstitution uniformly determines antiviral humoral function. Rather, antibody activity reflects the combined influence of host recovery, prior antigenic experience, and the degree of match between established immune memory and the virus being evaluated.

Within this broader relationship, the SARS-CoV-2 analyses further delineate how exposure history and subsequent viral divergence modify the serological consequences of CD4^+^ T-cell recovery. In both the initial and validation cohorts, BA.5 remained the predominant neutralization target across CD4 strata, whereas D614G activity varied more strongly with immune status. Antigenic cartography supported this distinction. The distance between BA.5 and D614G decreased with increasing CD4^+^ T-cell count, while the corresponding separation from BA.5 to NB.1.8.1 and XFG showed no comparable trend (**Figures 1 and 2**). Although the proportion of individuals for whom D614G represented the dominant target was highest among healthy controls, the overall distribution of variant-specific neutralization categories did not reach the conventional threshold for statistical significance (*p* = 0.057; Figure 2D). These data are therefore more consistent with a continuous shift in serological hierarchy than with discrete immune phenotypes. Reduced relative dominance of D614G in participants with lower CD4^+^ T-cell count was not accompanied by enhanced activity against NB.1.8.1, XFG, or more divergent sarbecoviruses. Attenuation of ancestral imprinting and expansion of neutralization breadth should therefore be regarded as distinct immunological phenomena. The population-level sequence of SARS-CoV-2 exposure in China offers an epidemiological basis for understanding this profile. Widespread use of ancestral-strain inactivated vaccines preceded large-scale BA.5.2/BF.7-associated infections and later circulation of additional Omicron lineages, resulting in repeated encounters with more recent viral antigens. Although this sequence cannot reconstruct individual exposure trajectories, it is compatible with the relative stability of BA.5 neutralization across CD4 strata and the stronger gradient observed for D614G. Previous studies have shown that memory B cells established by ancestral antigens can be preferentially recalled after Omicron infection, whereas repeated Omicron encounters can progressively reshape antibody specificity and epitope targeting, expand Omicron-specific B-cell clones, and broaden recognition of divergent variants. Antigenically updated vaccines can further enhance neutralization of matched and related viruses, although the extent of this redirection remains constrained by pre-existing ancestral imprinting.^11,22–24,30–34^

These observations suggest that host immune recovery may influence the relative serological expression of previously established memory rather than uniformly altering all virus-specific antibody responses. CD4^+^ T cells, particularly T follicular helper cells, contribute to germinal-center reactions, affinity maturation, class-switch recombination, and the establishment and maintenance of B-cell memory, providing biological plausibility for this interpretation.^35–37^ However, the present serological analyses cannot determine whether BA.5-dominant activity reflects de novo recruitment of B-cell clones, further maturation of cross-reactive memory generated by ancestral antigens, or contributions from both pathways. The imprinting described here should therefore be considered a serological phenotype rather than direct evidence of B-cell clonal origin or evolutionary trajectory. The attenuated CD4-associated gradient for NB.1.8.1 and XFG further suggests that the contribution of host recovery decreases as viral antigens become more distant from the pre-existing antibody repertoire. Under these conditions, immune escape may constrain neutralization more strongly than differences in CD4^+^ T-cell restoration. Host immune status and antigenic distance thus represent distinct determinants of antibody function. The former affects the maintenance and expression of established memory, whereas the latter limits its activity against divergent viral antigens.

The cross-pathogen analysis extends this interpretation beyond SARS-CoV-2. Neutralizing responses showed stronger within-pathogen correlations, whereas correlations among SARS-CoV-2, HIV-1, and influenza viruses were generally weak (**Figure 3C**). Multivariable analyses further revealed opposing relationships with CD4^+^ T-cell count. Higher count was linked to greater D614G neutralization but lower activity against CNE8, CH119.10, and H3N2. The interval from HIV-1 diagnosis to ART initiation was positively related to neutralization of all three HIV-1 strains, whereas longer ART duration corresponded to lower activity against CNE15 and CH119.10 (**Figure 3D**). The inverse relationship between HIV-1-specific activity and CD4^+^ T-cell levels may therefore reflect the combined influence of prior antigen burden, duration of untreated infection, viral diversification, and treatment course rather than enhanced humoral immunity in individuals with incomplete reconstitution. This view is consistent with previous studies linking HIV-1 neutralization breadth to infection duration, viral load, and viral diversity, and showing that suppression of ongoing antigenic stimulation after ART can alter antibody magnitude and persistence.^38–41^ The H3N2 finding may similarly reflect antibody memory accumulated through repeated infection, vaccination, and antigenic drift, whereas influenza B showed little dependence on CD4 status (**Figures 3B and 3D**). PLWH sera showed no detectable neutralization of SFTSV VSV pseudoviruses while retaining activity against HIV-1 pseudoviruses generated on the same VSV platform, arguing against residual antiretroviral drugs as an explanation for the HIV-1 findings (**Figure S2**). The contrasting behavior of these viruses reinforces the importance of pathogen-specific exposure trajectories in shaping the relationship between immune recovery and humoral function.

Neutralization-to-protection bridging provides a model-based perspective on the potential functional implications of these serological differences. Model-estimated protection against D614G increased from 43.8% in the CD4 <200 cells/µL group to 69.6% in the CD4 ≥500 cells/µL group and 79.9% in healthy controls. For BA.5, the corresponding values were higher across all cohorts, spanning 80.0%–89.5%. In contrast, estimates for NB.1.8.1 and XFG were lower, at 53.2%–59.1% and 51.1%–56.7%, respectively, with no monotonic trend across CD4^+^ T cell strata (**Figures 4B and 4C**). These differences are consistent with greater immune reconstitution being associated with stronger functional antibody activity when viral antigens remain relatively well matched to established memory, whereas extensive divergence may attenuate this relationship. Neutralizing antibody levels are quantitatively associated with protection from symptomatic SARS-CoV-2 infection,^17,42^ but the values reported here are conditional estimates derived from an external neutralization-to-protection model and a publicly available D614G reference. They should not be interpreted as observed infection rates, vaccine effectiveness, or individual probabilities of protection. The bridging analysis is therefore most informative for comparing relative functional profiles rather than assigning absolute clinical protection. Studies of XBB.1.5-, KP.2-, and more recent antigen-matched vaccines have shown that updating vaccine antigens can redirect serum neutralization and attenuate ancestral bias,^31,34,43^ further supporting the importance of correspondence between pre-existing immunity and circulating viruses. CD4^+^ T-cell count remains central to the assessment of immune reconstitution in PLWH, but they do not directly quantify functional antibody coverage against a specific pathogen or strain. Virus-specific serology may therefore complement conventional measures of immune recovery when considering booster vaccination and immune monitoring in immunocompromised populations.

Our study indicates that antiviral humoral immunity is shaped jointly by host immune recovery, prior antigenic experience, and subsequent viral evolution. The contribution of pre-existing memory depends both on its persistence within the host and on the extent to which later viral antigens remain recognizable by the established antibody repertoire. This relationship may account for the heterogeneous links between CD4^+^ T-cell status and antibody activity across pathogens. It also shows that attenuation of ancestral serological imprinting should not be equated with broader neutralization of divergent variants. CD4^+^ T-cell count therefore captures one aspect of immune status but do not directly define the breadth or effectiveness of virus-specific antibody responses. Functional serology may provide complementary information for assessing immune coverage in immunocompromised populations and for informing booster vaccination and immune monitoring.

## ACKNOWLEDGMENTS

This work was supported by the National Natural Science Foundation of China (grant no. 92569102 to L.L.). Support was also provided by the Taikang Center for Life and Medical Sciences, Wuhan University, Wuhan, China, and the State Key Laboratory of Virology and Biosafety, Wuhan University, Wuhan, China, to L.L. and Q.W., as well as by the 2026 National Science and Technology Major Program for New-Generation Artificial Intelligence (2026ZD0127500), including subprojects 2026ZD0127501 awarded to L.L. and 2026ZD0127503 awarded to Q.W. The funders had no role in study design, data collection and analysis, the decision to publish, or preparation of the manuscript.

## AUTHOR CONTRIBUTIONS

Q.W. and L.L. conceived and supervised the project. J.H. and S. Zhang. conducted experiments. S. Zou. and K.L. collected clinical samples. L.S. and T.L. helped with protein production for ELISA. C.X. and C.C. prepared SFTSV pseudovirus. C.L. performed the bioinformatic analyses. Q.W., L.L., J.H., S. Zhang., and C.L. analyzed the data and wrote the manuscript.

## RESOURCE AVAILABILITY

### Lead contact

Further information and requests for resources and reagents should be directed to and will be fulfilled by the lead contact, Prof. Lihong Liu.

### Materials availability

All requests for resources and reagents should be directed to and will be fulfilled by the Lead Contact, Prof. Lihong Liu. This includes cell lines, plasmids, antibodies, viruses, sera, and proteins. All reagents will be made available on request after completion of a Material Transfer Agreement.

### Data and code availability

Any additional information required to reanalyze the data reported in this paper is available from the lead contact upon request.

## EXPERIMENTAL MODEL AND SUBJECT DETAILS

### Clinical cohorts

Two independent serum cohorts were included in this study. The small exploratory cohort comprised 27 PLWH, including 11 with CD4^+^ T-cell count <200 cells/μL and 16 with count ≥500 cells/μL (**Table S1**). The validation cohort comprised 366 samples from 275 people living with HIV (PLWH) and 91 healthy controls (**Figure 2**). After excluding 11 PLWH with plasma HIV-1 RNA levels >200 copies/mL, the remaining 264 PLWH were stratified by CD4^+^ T-cell count into four groups: <200 (*n* = 28), 200 - 349 (*n* = 55), 350 - 499 (*n* = 70), and ≥500 cells/μL (*n* = 111), yielding a final analytical cohort of 355 participants.

All samples were collected between December 2025 and July 2026. Age, sex, time since HIV-1 diagnosis, CD4^+^ T-cell count, and current antiretroviral therapy (ART) regimen were recorded at serum collection. ART regimens were classified as integrase strand transfer inhibitors (INSTIs), non-nucleoside reverse transcriptase inhibitors (NNRTIs), protease inhibitors (PIs), or other regimens. Written informed consent was obtained from all participants, and the study was approved by the Ethics Committee of the Hubei Provincial Center for Disease Control and Prevention (Approval No. HBCDC-AF/SC-08/02.0).

### Cell lines

HEK293T (Cat# CRL-3216), Vero E6 (Cat# CRL-1586), and I1-Hybridoma (Cat# CRL-2700) cells were purchased from the American Type Culture Collection (ATCC). Expi293F (Cat# GDC0655) and Huh7.5.1 (Cat# GDC0137) cells were obtained from the China Center for Type Culture Collection (CCTCC). GHOST-CCR5-CXCR4 cells (GHOST (3) CXCR4+CCR5+; ARP-3942) were obtained from the NIH HIV Reagent Program, Division of AIDS, NIAID, NIH (contributed by Vineet N. KewalRamani and Dan R. Littman). The Vero E6-TMPRSS2-T2A-ACE2 cell line was generated by lentiviral-mediated gene transduction.

Adherent cells were routinely maintained at 37 °C with 5% CO₂ in Dulbecco’s modified Eagle medium (DMEM; Gibco, Cat# C11995500BT) supplemented with 10% fetal bovine serum (FBS; Umedium, Cat# 3023A) and 100 μg/mL penicillin-streptomycin. GHOST-CCR5-CXCR4 cells were maintained in DMEM supplemented with 10% FBS, 2 mM L-glutamine, 100 U/mL penicillin, 100 μg/mL streptomycin, 500 μg/mL G418, and 1 μg/mL puromycin. Expi293F cells were maintained in suspension in SMM 293-TII Expression Medium (Sino Biological, Cat# M293TII) containing penicillin-streptomycin at 37 °C with 5% CO₂ and continuous shaking at 125 rpm. I1-Hybridoma cells were maintained in Minimum Essential Medium (MEM; Gibco, Cat# C11095500BT) supplemented with 20% FBS and 100 μg/mL penicillin-streptomycin at 37 °C with 5% CO₂.

### Plasmids

Plasmids encoding coronavirus spike proteins, HIV-1 envelope proteins, TMPRSS11D, and influenza HA and NA were synthesized by GentleGen (Suzhou, China) and cloned into the pcDNA3.1 eukaryotic expression vector. To enhance pseudovirus infectivity, the cytoplasmic tail (CT) of each coronavirus spike was truncated. The spikes of D614G, B.1.1.7, B.1.351, B.1.617.2, BA.1, BA.2, BA.5, BF.7, XBB.1.5, EG.5.1, BA.2.86, JN.1, KP.3.1.1, XEC, LP.8.1, LF.7, NB.1.8.1, XFG, PE.1.4, PY.1.1.1, BA.3.2.1, and BA.3.2.2 were truncated by deletion of the C-terminal 18 amino acids; those of GD-Pangolin and GX-Pangolin by deletion of the C-terminal 17 amino acids; and those of SARS-CoV-1 and WIV1 by deletion of the C-terminal 19 and 13 amino acids, respectively. For the HIV-1 envelope constructs, the native transmembrane domain (TM) and CT were replaced with the corresponding regions of the vesicular stomatitis virus glycoprotein (VSV-G) to enhance envelope-mediated pseudovirus incorporation and infectivity.

### Pseudovirus production

Pseudoviruses were generated as previously described^20,21^. In brief, HEK293T cells were seeded in 15-cm culture dishes and, on the following day, were transfected with 30 µg of plasmids encoding coronavirus spike or chimeric HIV-1 envelope using linear polyethylenimine (PEI, YEASEN, Cat# 40816ES03) at a DNA:PEI ratio of 1:3. For influenza pseudovirus production, plasmids encoding HA, NA, and TMPRSS11D were co-transfected at a 3:1:2 ratio. At 24 h post-transfection, cells were infected with VSV-G-pseudotyped ΔG-luciferase rVSV (Kerafast, Cat# EH1020-PM). Two hours after infection, the inoculum was removed, cells were washed twice with DMEM containing 2% FBS and 1% penicillin-streptomycin, and fresh culture medium was added. At 24 h post-infection, culture supernatants containing pseudoviruses were harvested and clarified by centrifugation at 2,000 rpm for 5 min. To eliminate residual VSV-G-pseudotyped input virus, clarified supernatants were incubated with I1-Hybridoma supernatant. Pseudovirus preparations were aliquoted and stored at −80 °C until use.

### Pseudovirus neutralization assay

Pseudoviruses were titrated before each neutralization assay to normalize the viral input between assays. Serum samples were heat-inactivated at 56 °C for 30 min before use. For determination of the 50% inhibitory dilution (ID_50_), sera were serially diluted fourfold starting at a 1:20 dilution. Equal volumes (50 µL) of diluted serum and pseudovirus were mixed and incubated at 37 °C for 1 h. Target cells were then added at 100 µL per well, and the plates were incubated overnight at 37 °C in a humidified 5% CO₂ incubator. Luciferase activity was measured using a Varioskan LUX multimode microplate reader (Thermo Fisher Scientific). The ID_50_ was defined as the serum dilution at which relative luciferase units were reduced by 50% relative to virus-only control wells.

### Protein purification

Recombinant spike proteins of SARS-CoV-2 D614G, BA.5, NB.1.8.1, and XFG were produced in Expi293F cells (Figure S3A). When cell density reached 2.5 × 10⁶ cells/mL, cells were transfected with the corresponding spike expression plasmids using PEI. Five days post-transfection, culture supernatants were collected and clarified by centrifugation at 8,000 rpm for 20 min at 4 °C. The supernatant pH was adjusted by adding 1/20 volume of 1 M Tris-HCl (pH 8.0), followed by thorough mixing, and the supernatants were filtered through a 0.45 µm membrane. His-tagged spike proteins were captured by passing the filtrate through protein purification resin (YEASEN, Cat# 20497ES) four times. The resin was washed sequentially with 20 mM and 50 mM imidazole, and bound proteins were eluted with 250 mM imidazole. Eluted fractions were buffer-exchanged into 1×PBS by ultrafiltration, and protein concentrations were determined after buffer exchange.

### Enzyme-linked immunosorbent assay (ELISA)

Binding of serum antibodies to purified SARS-CoV-2 spike proteins (D614G, BA.5, NB.1.8.1, and XFG) was measured by ELISA. Spike proteins were diluted in phosphate-buffered saline (PBS) and coated onto 96-well plates at 50 ng per well overnight at 4 °C. The following day, the coating solution was removed, wells were washed twice with PBST (PBS containing 0.1% Tween-20), and plates were blocked with 200 µL blocking buffer (PBST supplemented with 1% bovine serum albumin [BSA; Solarbio, Cat# A8020] and 1% FBS) at 37 °C for 1 h. After two washes with PBST, serum samples were serially diluted fourfold in blocking buffer starting at a 1:100 dilution, and 100 µL of each dilution was added to duplicate wells and incubated at 37 °C for 1 h. Plates were washed three times with PBST and incubated with 100 µL per well of horseradish peroxidase (HRP)-conjugated goat anti-human IgG (H+L) (Jackson ImmunoResearch, Cat# 109-035-088; 1:10,000 in blocking buffer) at 37 °C for 1 h, followed by four washes with PBST. The reaction was developed with 100 µL per well of TMB substrate (Solarbio, Cat# PR1200) for 5 min at room temperature in the dark, stopped with 50 µL of 2 M H₂SO₄, and absorbance at 450 nm was recorded on a Varioskan LUX microplate reader. All samples were tested in duplicate, and the mean OD450 of duplicate wells was used for subsequent analysis.

### Antigenic cartography

Antigenic maps were generated by integrating individual-serum ID_50_ values in two dimensions^44^ using Racmacs (version 1.2.9; https://acorg.github.io/Racmacs/) in R version 4.6.1. Maps were optimized using 2,000 optimizations with the minimum-column-basis parameter set to none and with disconnected points ignored. Sera contributing fewer than two measurements or fewer than two distinct ID_50_ values were excluded, and each group map was fitted from at least ten usable sera. Antigenic distances were calculated as Euclidean distances between optimized variant coordinates. For the expanded SARS-CoV-2 map in **Figure 1C**, the optimized configuration was displayed without post-hoc re-centering on a specific variant. Two predominantly assay-floor variants (GX-Pangolin and WIV1) were excluded before map fitting. For the four-variant SARS-CoV-2 maps in **Figure 2**, D614G, BA.5, NB.1.8.1, and XFG were mapped separately within each clinical group. Each configuration was translated to place BA.5 at the origin, then rotated and, when required, reflected to position D614G to the left of NB.1.8.1 and XFG. Maps were displayed on fixed antigenic-unit grids. The y-axis was compressed by 0.76 for figure layout only; numerical BA.5-to-variant distance labels retained the original coordinate-based distances.

### Dominant neutralization response composition

For sera with complete positive ID_50_ measurements against D614G, BA.5, NB.1.8.1, and XFG, the dominant variant was defined directly from the untransformed titers as the variant with the unique highest ID_50_ value:

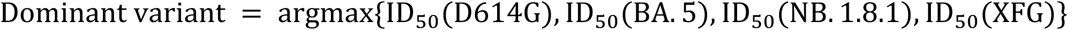

Thus, each serum was assigned to the variant with the highest ID_50_. This rank-based definition is equivalent to identifying the variant with a relative deficit of zero, but the composition analysis was calculated directly from the raw ID_50_ matrix rather than from a delta variable.

Sera with all four responses at or below the assay floor (ID_50_ ≤20; *n*=5) were excluded because a dominant variant could not be determined. The analysis therefore included 350 sera (27, 54, 69, 109, and 91 in the four CD4^+^ T-cell count strata and healthy controls, respectively). Group-specific dominant-variant compositions were displayed as pie charts. The overall distribution across the five groups was compared using Fisher’s exact test with Monte Carlo simulation (199,999 replicates; *p* = 0.057); this single omnibus test was not adjusted for multiple comparisons.

### Multivariable regression analysis

For each of the nine viruses, a separate linear regression model was fitted among PLWH with log_10_(ID_50_) as the outcome. Continuous predictors were standardized to z scores, where z(x) denotes (x − mean(x))/SD(x). ART class was coded as INSTI-based versus non-INSTI, and I (INSTI-based ART) is an indicator variable coded 1 for an INSTI-based regimen and 0 otherwise. Sex was not included because of marked sex imbalance among PLWH. The fitted model was:

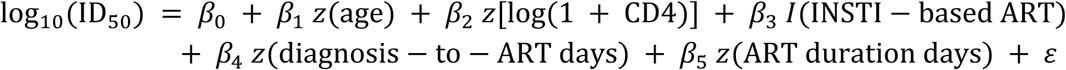

Regression coefficients and 95% confidence intervals were exponentiated as 10^β^ to report adjusted fold changes in ID_50_. Regression estimates were not adjusted for multiple comparisons and were interpreted separately for each virus. All tests were two-sided, and *P*<0.05 was considered statistically significant. Figures were generated in R using ggplot2 (version 4.0.2).

### Immune-bridging analysis

Public serum/plasma neutralization records were curated from the Stanford COVID-DRDB database (https://covdb.stanford.edu/; raw data payload: https://github.com/hivdb/covid-drdb-payload), restricted to records derived from Chinese serum or plasma samples, and from PubMed-indexed literature (https://pubmed.ncbi.nlm.nih.gov/). Literature curation focused on Chinese serum or plasma neutralization studies reporting prototype/D614G or contemporary Omicron-variant neutralizing antibody titers. In total, 111 cohort-level neutralization entries from 67 published studies were retained (83 entries covering 57 studies from the database and 28 entries extracted from 11 additional publications; 33 studies contributed more than one entry), representing vaccinee (*n* = 56), convalescent (*n* = 35), and vaccinated-plus-infected (*n* = 20) cohorts, with no PLWH cohorts represented; the entries comprised 7,987 participants, and three entries did not report subgroup sizes. All titers represent 50% neutralization titers expressed as reciprocal dilutions (labeled NT50 in 107 entries, of which one used the synonym NAT50, ID_50_ in one entry, and IC_50_ in three entries) and were analyzed as dimensionless values; 93 entries used pseudovirus-based and 18 authentic-virus assays. PMID-indexed references were retained in the supplementary reference appendix.

For each entry, the public dataset documents how the D614G value was established: 41 entries reported an explicit D614G arm; 41 entries reported only an ancestral/WT strain (Wuhan-Hu-1, WA1/2020, HKU-001a, VM20001061-2, JS02, ZJ-WGF, and related lineage A/B viruses carrying the ancestral D614), which was used as a pre-specified surrogate for D614G. For each clinical group and variant, serum neutralization titers from the study cohort were divided by this anchor to obtain normalized neutralization levels. Normalized values were mapped to protection against symptomatic SARS-CoV-2 infection using the neutralization–protection model of Khoury et al.^17^:

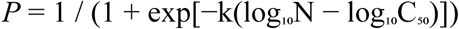

where N is the neutralization titer relative to the public D614G anchor, k = 2.543306 is the Hill slope of the logistic relationship, and C_50_ = 0.2118421 is the relative neutralization level at which 50% protection is predicted. Group-level mean protection and 95% confidence intervals were estimated using 2,000 bootstrap resamples of individual normalized titers within each group and variant. The resulting values are model-based immune-bridging estimates and do not represent observed infection rates, vaccine effectiveness, or individual clinical protection.

## QUANTIFICATION AND STATISTICAL ANALYSIS

Geometric mean titers (GMTs) and 95% confidence intervals were calculated on the log_10_(ID_50_) scale and back-transformed for display. Individual ID_50_ values were plotted on a log_10_ axis. Differences between the five clinical groups were assessed using two-sided Wilcoxon rank-sum tests; all 10 pairwise comparisons were performed for each of the nine viruses, and *P* values were adjusted within virus using the Benjamini–Hochberg (BH) method. Pairwise virus correlations were evaluated separately in PLWH and healthy controls using Spearman’s rank correlation, followed by BH adjustment within each population.

Clinical correlations of age, CD4^+^ T-cell count, ART duration, and diagnosis-to-ART duration with raw ID_50_ values were assessed among PLWH using Spearman’s rank correlation, with BH adjustment across 36 tests (four clinical variables and nine viruses). Viral load was not included in the final clinical heatmap. Because log_10_ transformation preserves the ranks of positive ID_50_ values, the corresponding raw-ID_50_ and log_10_(ID_50_) analyses would yield identical correlation coefficients and *P* values when based on the same observations.

