## Supplemental figures and tables for "Immune Reconstitution and Antigenic Exposure History Shape Virus-Specific Humoral Immunity in People with HIV"

### SUPPLEMENTAL INFORMATION

**Table S1. Clinical characteristics of 27 people living with HIV (PLWH) with viral load ≤200 copies/mL, related to Figure 1.** PLWH were stratified by peripheral $CD4^+$ T-cell count into <200 (*n*=11) and ≥500 (*n*=16) cells/µL. Serum samples from these two small cohorts were tested against 26 coronaviruses in **Figure 1**.

| Characteristics | $CD4^+$ T-cell count among PLWH (n=27) | |
|---|---|---|
| | **<200 (n=11)** | **≥500 (n=16)** |
| Male sex - no. (%) | 11 (100) | 14 (87.5) |
| Mean age - years | 47 | 43 |
| Mean days since HIV-1 diagnosis | 1174 | 2112 |
| Mean $CD4^+$ T-cell count - cells/µL | 153 | 753 |
| Current ART regimen - no. (%) | | |
| Integrase strand transfer inhibitor | 9 (81.8) | 15 (93.8) |
| Non-nucleoside reverse transcriptase inhibitor | 2 (18.2) | 1 (6.2) |

**Table S2. Public neutralization dataset used for neutralization-to-protection bridging (111 cohort-level entries from 67 published studies of Chinese serum or plasma samples)**, related to Figure 4**.** Each entry documents the source, PMID, study, cohort category, number of subjects, platform, and GMTs for D614G, BA.5, NB.1.8.1, and XFG. GMTs are expressed as reciprocal dilution titers; — indicates not tested.

| No. | PMID | Study | Source | Cohort category | n subjects | Platform | D614G GMT | BA.5 GMT | NB.1.8.1 GMT | XFG GMT |
|---|---|---|---|---|---|---|---|---|---|---|
| 1 | 34021265 | Cao 2021 | Cov-RDB | Vaccinee | 40 | pseudovirus | 406.2 | — | — | — |
| 2 | 34484190 | Ding 2021 | Cov-RDB | Convalescent | 5 | pseudovirus | 2017 | — | — | — |
| 3 | 33633321 | Hu 2021 | Cov-RDB | Convalescent | 40 | pseudovirus | 316.7 | — | — | — |
| 4 | 33870240 | Huang 2021 | Cov-RDB | Vaccinee | 24 | pseudovirus | 99.9 | — | — | — |
| 5 | 33735608 | Li 2021 | Cov-RDB | Convalescent | 18 | pseudovirus | 1303 | — | — | — |
| 6 | 33580167 | Li 2021 | Cov-RDB | Convalescent | 48 | pseudovirus | 980.2 | — | — | — |
| 7 | 34166623 | Wang 2021 | Cov-RDB | Convalescent | 24 | pseudovirus | 772 | — | — | — |
| 8 | 33822491 | Wang 2021 | Cov-RDB | Convalescent | 34 | pseudovirus | 167.9 | — | — | — |
| 9 | 33822491 | Wang 2021 | Cov-RDB | Vaccinee | 50 | pseudovirus | 53.6 | — | — | — |
| 10 | 34704423 | Xie 2021 | Cov-RDB | Vaccinee | 8 | pseudovirus | 101.6 | — | — | — |
| 11 | 34649620 | Yi 2021 | Cov-RDB | Convalescent | 9 | pseudovirus | 2931.1 | — | — | — |
| 12 | 34458880 | Zhao 2021 | Cov-RDB | Vaccinee | 28 | pseudovirus | 1103.6 | — | — | — |
| 13 | 34935594 | Ai 2022 | Cov-RDB | Vaccinated + infected | 7 | pseudovirus | 1794.4 | — | — | — |
| 14 | 34935594 | Ai 2022 | Cov-RDB | Vaccinee | 70 | pseudovirus | 177.7 | — | — | — |
| 15 | 35594867 | Ai 2022 | Cov-RDB | Vaccinee | 58 | pseudovirus | 327.9 | — | — | — |
| 16 | 34862467 | Cao 2022 | Literature | Vaccinee | 41 | authentic virus | 794 | — | — | — |
| 17 | 34862467 | Cao 2022 | Literature | Vaccinee | 81 | authentic virus | 1306 | — | — | — |
| 18 | 35714668 | Cao 2022 | Cov-RDB | Convalescent | 78 | pseudovirus | 1153.8 | 54.9 | — | — |
| 19 | 35714668 | Cao 2022 | Cov-RDB | Vaccinee | 78 | pseudovirus | 736 | 87.9 | — | — |
| 20 | 36270286 | Cao 2022 | Cov-RDB | Vaccinated + infected | 95 | pseudovirus | 1309 | 174.7 | — | — |
| 21 | 36270286 | Cao 2022 | Cov-RDB | Vaccinee | 40 | pseudovirus | 632 | 75 | — | — |
| 22 | 35084295 | Chen 2022 | Cov-RDB | Convalescent | 15 | pseudovirus | 127 | — | — | — |
| 23 | 35084295 | Chen 2022 | Cov-RDB | Vaccinee | 55 | pseudovirus | 140.4 | — | — | — |
| 24 | 35608053 | Chen 2022 | Cov-RDB | Vaccinated + infected | 10 | pseudovirus | 3581 | — | — | — |
| 25 | 35608053 | Chen 2022 | Cov-RDB | Vaccinee | 154 | pseudovirus | 85.5 | — | — | — |
| 26 | 35051989 | Cheng 2022 | Cov-RDB | Convalescent | 88 | pseudovirus | 186.8 | — | — | — |
| 27 | 35051989 | Cheng 2022 | Cov-RDB | Vaccinee | 146 | pseudovirus | 135.2 | — | — | — |
| 28 | 35514306 | Cheng 2022 | Cov-RDB | Convalescent | 27 | pseudovirus | 9.2 | — | — | — |
| 29 | 35514306 | Cheng 2022 | Cov-RDB | Vaccinated + infected | 73 | pseudovirus | 217.5 | — | — | — |
| 30 | 35514306 | Cheng 2022 | Cov-RDB | Vaccinee | 60 | pseudovirus | 185.9 | — | — | — |
| 31 | 36399969 | Chuang 2022 | Cov-RDB | Vaccinee | 240 | pseudovirus | 618 | — | — | — |
| 32 | 35966041 | Duan 2022 | Cov-RDB | Convalescent | 342 | pseudovirus | 170.4 | — | — | — |
| 33 | 35966041 | Duan 2022 | Cov-RDB | Vaccinee | 106 | pseudovirus | 51.4 | — | — | — |
| 34 | 34877393 | Hu 2022 | Cov-RDB | Convalescent | 20 | pseudovirus | 115.7 | — | — | — |
| 35 | 34877393 | Hu 2022 | Cov-RDB | Vaccinee | 30 | pseudovirus | 103.6 | — | — | — |
| 36 | 35017716 | Hu 2022 | Cov-RDB | Convalescent | 12 | pseudovirus | 295 | — | — | — |
| 37 | 35017716 | Hu 2022 | Cov-RDB | Vaccinee | 12 | pseudovirus | 216 | — | — | — |
| 38 | 36179744 | Jian 2022 | Cov-RDB | Convalescent | 97 | pseudovirus | 1391.6 | 134.7 | — | — |
| 39 | 36179744 | Jian 2022 | Cov-RDB | Vaccinee | 40 | pseudovirus | 652 | 53 | — | — |
| 40 | 35791773 | Jiang 2022 | Cov-RDB | Vaccinee | 210 | pseudovirus | 172.9 | — | — | — |
| 41 | 35260792 | Ju 2022 | Cov-RDB | Convalescent | 19 | pseudovirus | 2507.7 | — | — | — |
| 42 | 36560521 | Li 2022 | Cov-RDB | Vaccinee | 129 | pseudovirus | 6.4 | — | — | — |
| 43 | 34739037 | Lien 2022 | Cov-RDB | Vaccinee | 45 | pseudovirus | 246.9 | — | — | — |
| 44 | 34746904 | Liu 2022 | Cov-RDB | Convalescent | 16 | pseudovirus | 286.4 | — | — | — |
| 45 | 34915551 | Lu 2022 | Cov-RDB | Vaccinee | 50 | pseudovirus | 70.6 | — | — | — |
| 46 | 36338432 | Peng 2022 | Cov-RDB | Convalescent | 53 | pseudovirus | 1115 | — | — | — |
| 47 | 35266815 | Ren 2022 | Cov-RDB | Convalescent | 28 | pseudovirus | 1317.8 | — | — | — |
| 48 | 35034583 | Wang 2022 | Cov-RDB | Convalescent | 10 | pseudovirus | 1105.8 | — | — | — |
| 49 | 35034583 | Wang 2022 | Cov-RDB | Vaccinee | 30 | pseudovirus | 426.3 | — | — | — |
| 50 | 35090164 | Wang 2022 | Cov-RDB | Vaccinee | 120 | pseudovirus | 92.2 | — | — | — |
| 51 | 35001836 | Wang 2022 | Cov-RDB | Convalescent | 16 | pseudovirus | 1328 | — | — | — |
| 52 | 35001836 | Wang 2022 | Cov-RDB | Vaccinee | 20 | pseudovirus | 670 | — | — | — |

| No. | PMID | Study | Source | Cohort category | n subjects | Platform | D614G GMT | BA.5 GMT | NB.1.8.1 GMT | XFG GMT |
|---|---|---|---|---|---|---|---|---|---|---|
| 53 | 36229428 | Wang 2022 | Cov-RDB | Convalescent | 12 | pseudovirus | 3130 | 467 | — | — |
| 54 | 36229428 | Wang 2022 | Cov-RDB | Vaccinee | 20 | pseudovirus | 572.5 | 52.3 | — | — |
| 55 | 35034952 | Yu 2022 | Cov-RDB | Vaccinee | 292 | pseudovirus | 294.9 | — | — | — |
| 56 | 35176972 | Yu 2022 | Cov-RDB | Vaccinee | 200 | pseudovirus | 239 | — | — | — |
| 57 | 35401825 | Zhang 2022 | Cov-RDB | Convalescent | 48 | pseudovirus | 55 | — | — | — |
| 58 | 35401825 | Zhang 2022 | Cov-RDB | Vaccinee | 60 | pseudovirus | 27.5 | — | — | — |
| 59 | 35379815 | Zhang 2022 | Cov-RDB | Convalescent | 16 | pseudovirus | 404.1 | — | — | — |
| 60 | 35379815 | Zhang 2022 | Cov-RDB | Convalescent | 690 | pseudovirus | 58.2 | — | — | — |
| 61 | 35793198 | Zhao 2022 | Cov-RDB | Vaccinee | 40 | pseudovirus | 630.1 | 30.8 | — | — |
| 62 | 35793198 | Zhao 2022 | Cov-RDB | Vaccinee | 154 | pseudovirus | 460.9 | 27.8 | — | — |
| 63 | 35592756 | Zhou 2022 | Cov-RDB | Vaccinee | 63 | pseudovirus | 301.4 | — | — | — |
| 64 | 35429398 | Zhu 2022 | Cov-RDB | Convalescent | 36 | pseudovirus | 563 | — | — | — |
| 65 | 35429398 | Zhu 2022 | Cov-RDB | Vaccinee | 31 | pseudovirus | 3942 | — | — | — |
| 66 | 36309305 | Zhu 2022 | Cov-RDB | Convalescent | 20 | pseudovirus | 4914.1 | — | — | — |
| 67 | 36309305 | Zhu 2022 | Cov-RDB | Vaccinee | 30 | pseudovirus | 1709.7 | — | — | — |
| 68 | 36535326 | Cao 2023 | Cov-RDB | Vaccinated + infected | 110 | pseudovirus | 1318.7 | 65.1 | — | — |
| 69 | 36535326 | Cao 2023 | Cov-RDB | Vaccinee | 40 | pseudovirus | 652 | 33 | — | — |
| 70 | 36946504 | Cheung 2023 | Cov-RDB | Vaccinated + infected | 39 | pseudovirus | 941.2 | 80.7 | — | — |
| 71 | 36946504 | Cheung 2023 | Cov-RDB | Vaccinee | 24 | pseudovirus | 345.4 | 4.4 | — | — |
| 72 | 36108092 | Cowling 2023 | Cov-RDB | Vaccinated + infected | 42 | pseudovirus | 320 | — | — | — |
| 73 | 36108092 | Cowling 2023 | Cov-RDB | Vaccinee | 138 | pseudovirus | 99.2 | — | — | — |
| 74 | 35675370 | Leung 2023 | Cov-RDB | Vaccinee | 40 | pseudovirus | 60.6 | — | — | — |
| 75 | 35675370 | Leung 2023 | Cov-RDB | Vaccinee | 40 | pseudovirus | 47.6 | — | — | — |
| 76 | 37549680 | Leung 2023 | Cov-RDB | Vaccinee | 80 | pseudovirus | 293.4 | — | — | — |
| 77 | 37313604 | Liu 2023 | Literature | Vaccinated + infected | 63 | pseudovirus | 742 | 533 | — | — |
| 78 | 36680413 | Shen 2023 | Cov-RDB | Convalescent | 13 | pseudovirus | 1095.7 | 97.6 | — | — |
| 79 | 36680413 | Shen 2023 | Cov-RDB | Vaccinated + infected | 54 | pseudovirus | 1481.9 | 197.2 | — | — |
| 80 | 37400857 | Wang 2023 | Literature | Convalescent | 15 | authentic virus | 125.1 | 23 | — | — |
| 81 | 37400857 | Wang 2023 | Literature | Vaccinated + infected | 24 | authentic virus | 555.9 | 41 | — | — |
| 82 | 37400857 | Wang 2023 | Literature | Vaccinee | 101 | authentic virus | 589.3 | 21.6 | — | — |
| 83 | 37400857 | Wang 2023 | Literature | Vaccinee | 12 | authentic virus | 1128.3 | 43 | — | — |
| 84 | 37400857 | Wang 2023 | Literature | Vaccinee | 23 | authentic virus | 1191.2 | 52.4 | — | — |
| 85 | 36519597 | Wang 2023 | Cov-RDB | Vaccinee | 23 | pseudovirus | 166.3 | — | — | — |
| 86 | 37369648 | Wang 2023 | Cov-RDB | Vaccinated + infected | 68 | pseudovirus | 1744.5 | 398.5 | — | — |
| 87 | 37369648 | Wang 2023 | Cov-RDB | Vaccinee | 22 | pseudovirus | 611.2 | 30 | — | — |
| 88 | 37336889 | Yang 2023 | Literature | Vaccinated + infected | 108 | pseudovirus | 392 | 317 | — | — |
| 89 | 36746173 | Yue 2023 | Cov-RDB | Vaccinated + infected | 126 | pseudovirus | 1435.4 | — | — | — |
| 90 | 36549416 | Zhang 2023 | Cov-RDB | Vaccinee | 451 | pseudovirus | 22 | — | — | — |
| 91 | 38801319 | Jia 2024 | Literature | Vaccinee | — | authentic virus | 177 | — | — | — |
| 92 | 40078728 | Liu 2024 | Literature | Vaccinee | 289 | pseudovirus | 27.6 | — | — | — |
| 93 | 40078728 | Liu 2024 | Literature | Vaccinee | 286 | pseudovirus | 127.1 | — | — | — |
| 94 | 39287459 | Wu 2024 | Literature | Convalescent | 137 | pseudovirus | 819.5 | — | — | — |
| 95 | 37993710 | Yisimayi 2024 | Cov-RDB | Vaccinated + infected | 291 | pseudovirus | 397.1 | 357.6 | — | — |
| 96 | 38191194 | Yuan 2024 | Literature | Vaccinee | 17 | authentic virus | 72 | 20 | — | — |
| 97 | 38191194 | Yuan 2024 | Literature | Vaccinee | 68 | authentic virus | 57 | 12 | — | — |
| 98 | 38191194 | Yuan 2024 | Literature | Vaccinee | 23 | authentic virus | 248 | 40 | — | — |
| 99 | 39568037 | Zhang 2024 | Literature | Convalescent | 107 | authentic virus | 280 | 41 | — | — |
| 100 | 39568037 | Zhang 2024 | Literature | Convalescent | 4 | authentic virus | 152 | 38 | — | — |
| 101 | 39568037 | Zhang 2024 | Literature | Convalescent | 91 | authentic virus | 287 | 45 | — | — |
| 102 | 39568037 | Zhang 2024 | Literature | Convalescent | 12 | authentic virus | 287 | 23 | — | — |
| 103 | 39568037 | Zhang 2024 | Literature | Convalescent | 10 | authentic virus | 362 | 119 | — | — |
| 104 | 36719034 | Chen 2023 | Cov-RDB | Vaccinated + infected | 76 | pseudovirus | — | 42.3 | — | — |
| 105 | 36591327 | Zhou 2023 | Literature | Vaccinated + infected | 5 | pseudovirus | — | 957 | — | — |
| 106 | 36591327 | Zhou 2023 | Literature | Vaccinated + infected | 3 | pseudovirus | — | 200 | — | — |
| 107 | 36591327 | Zhou 2023 | Literature | Vaccinated + infected | 1 | pseudovirus | — | 94 | — | — |
| 108 | 40078728 | Liu 2024 | Literature | Vaccinee | — | authentic virus | — | 8.8 | — | — |
| 109 | 40078728 | Liu 2024 | Literature | Vaccinee | — | authentic virus | — | 11.7 | — | — |
| 110 | 40484018 | Guo 2025 | Literature | Vaccinated + infected | 29 | pseudovirus | — | — | 76 | 67 |
| 111 | 40484018 | Guo 2025 | Literature | Vaccinated + infected | 21 | pseudovirus | — | — | 84 | 66 |

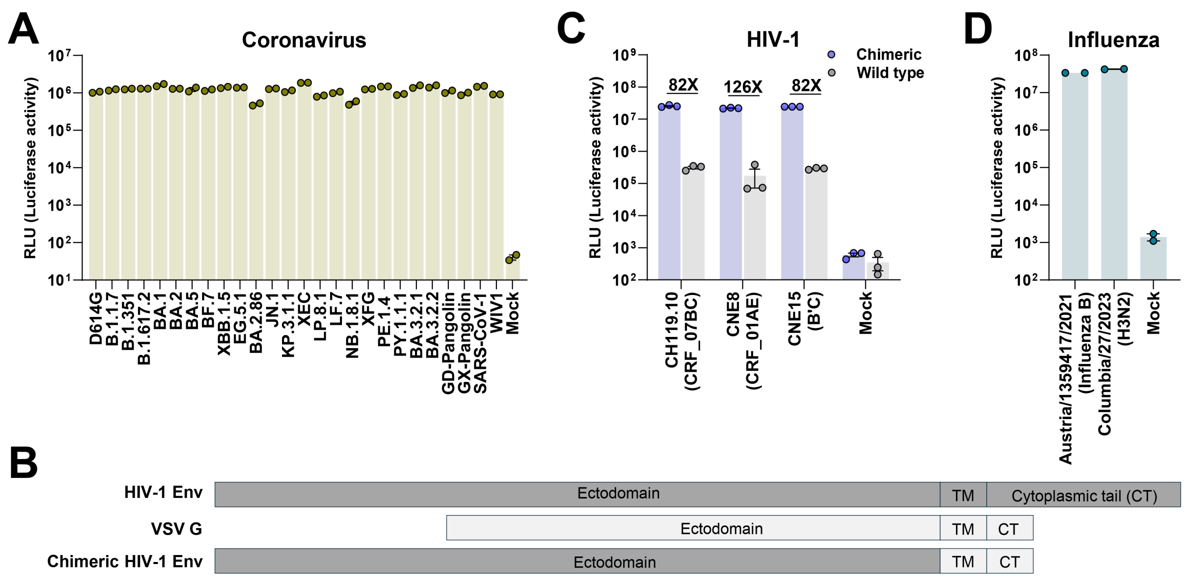


**Figure S1. Generation and infectivity of VSV pseudoviruses bearing different viral glycoproteins, related to Figures 1 and 3.**

**A.** Infectivity of VSV pseudoviruses bearing the indicated SARS-CoV-2 variant spikes and other related sarbecovirus spikes. SARS-CoV-2, SARS-CoV-1, and GD-Pangolin were tested in Vero E6 cells, whereas the GX-Pangolin and WIV1 pseudoviruses were tested in Vero E6-TMPRSS2-T2A-ACE2 cells. Infectivity was quantified by luciferase activity and expressed as relative luminescence units (RLU).

**B.** Schematic of the chimeric HIV-1 envelope design. Wild-type HIV-1 envelope comprises an ectodomain, transmembrane domain (TM), and cytoplasmic tail (CT). In the chimeric construct, the native HIV-1 envelope TM and CT were replaced with the corresponding regions of vesicular stomatitis virus glycoprotein (VSV-G), while the HIV-1 envelope ectodomain was retained.

**C.** Infectivity of VSV pseudoviruses bearing chimeric or wild-type HIV-1 envelope in GHOST-CCR5-CXCR4 cells. Fold differences in infectivity between chimeric envelope-bearing and the corresponding wild-type envelope-bearing pseudoviruses are shown above the bars.

**D.** Infectivity of VSV pseudoviruses bearing hemagglutinin (HA) and neuraminidase (NA) from B/Austria/1359417/2021 (influenza B) or A/Columbia/27/2023 (H3N2) in Huh7.5.1 cells.

For panels **A**, **C**, and **D**, data are shown as mean ± SEM. Mock denotes cells without pseudovirus infection.

**A**

| Characteristics | CD4+ T-cell count among PLWH (n=45) | | | |
|---|---|---|---|---|
| | < 200 (n=9) | 200 - 349 (n=12) | 350 - 499 (n=12) | ≥ 500 (n=12) |
| Male sex - no. (%) | 7 (77.8) | 12 (100) | 10 (83.3) | 12 (100) |
| Mean age - years | 57 | 41 | 36 | 30 |
| Mean days since HIV-1 diagnosis | 1356 | 1687 | 1235 | 1834 |
| Mean CD4+ T-cell count - cells/μL | 164 | 286 | 405 | 647 |
| Current ART regimen - no. (%) | | | | |
| Integrase strand transfer inhibitor | | | | |
| Biktarvy | 4 (44.4) | 4 (33.3) | 4 (33.3) | 4 (33.3) |
| Dovato | 3 (33.3) | 4 (33.3) | 4 (33.3) | 4 (33.3) |
| Non-nucleoside reverse transcriptase inhibitor | | | | |
| Efavirenz | 2 (22.2) | 4 (33.3) | 4 (33.3) | 4 (33.3) |

**B**

| Characteristics | SFTSV-infected individuals (n=9) |
|---|---|
| Male sex - no. (%) | 5 (55.6) |
| Female sex - no. (%) | 4 (44.4) |
| Mean age - years | 70 |
| Time to symptom onset - days | 15 |

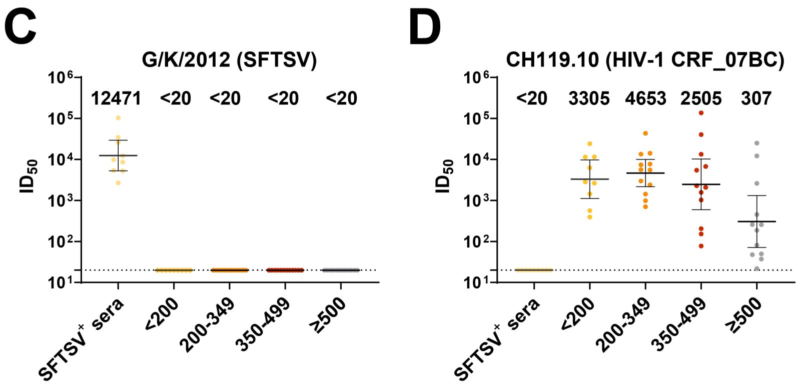


**Figure S2. Assessment of potential interference by antiretroviral therapy in VSV pseudovirus neutralization assays, related to Figure 1.**

**A.** Clinical characteristics of selected donors from four PLWH groups stratified by $CD4^+$ T-cell count (<200, 200–349, 350–499, and ≥500 cells/μL).

**B.** Clinical characteristics of the SFTSV-infected control cohort ($n$ = 9). SFTSV, severe fever with thrombocytopenia syndrome virus.

**C.** Serum neutralization of VSV pseudovirus bearing the Gn/Gc glycoproteins of SFTSV strain G/K/2012. Sera from individuals with prior SFTSV infection showed potent neutralizing activity, whereas sera from ART-treated PLWH showed no detectable neutralization, supporting the absence of nonspecific inhibition of the VSV pseudovirus by residual antiretroviral drugs.

**D.** Serum neutralization of VSV pseudovirus bearing HIV-1 CH119.10 envelope. Sera from PLWH neutralized the HIV-1 pseudovirus, whereas sera from SFTSV-infected individuals showed no detectable neutralization.

In panels **C** and **D**, points represent individual serum samples; horizontal lines and error bars indicate GMTs and 95% CIs. The dashed horizontal line denotes the assay detection limit ($ID_{50}$ = 20), and GMTs are shown above the plots.

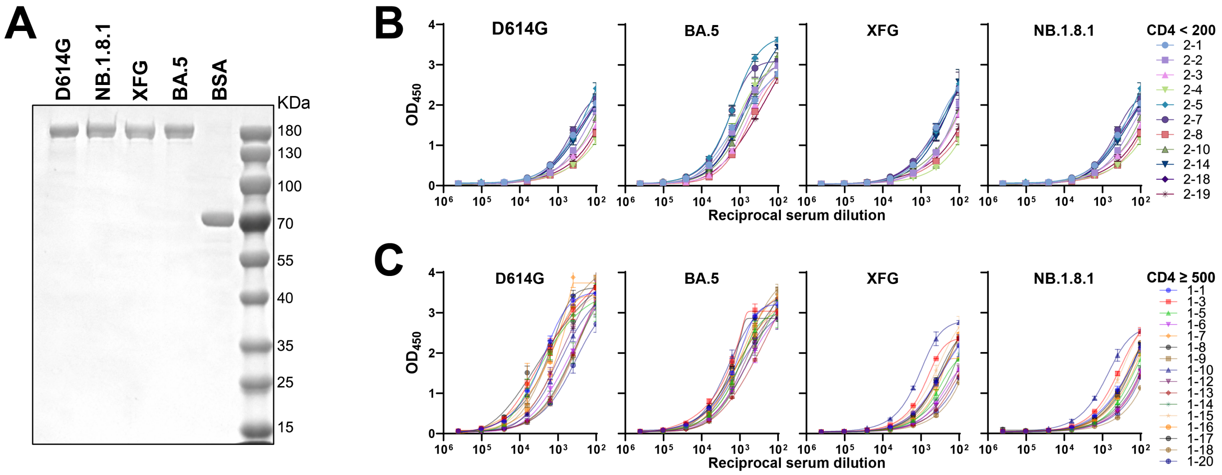


**Figure S3. Characterization of SARS-CoV-2 spikes and serum antibody binding in PLWH with different CD4$^{+}$ T-cell counts, related to Figure 1.**

**A.** Coomassie blue staining of SDS-PAGE gels showing purified spikes of SARS-CoV-2 D614G, BA.5, NB.1.8.1, and XFG.

**B, C**. Serum antibody binding to the indicated SARS-CoV-2 spikes was measured by ELISA in PLWH with CD4$^{+}$ T-cell count <200 cells/μL (**B**, *n* = 11) or ≥500 cells/μL (**C**, *n* = 16). Serum samples were serially diluted, and antibody binding was assessed by measuring optical density at 450 nm (OD450). Binding curves for the indicated spike proteins are shown for each cohort.

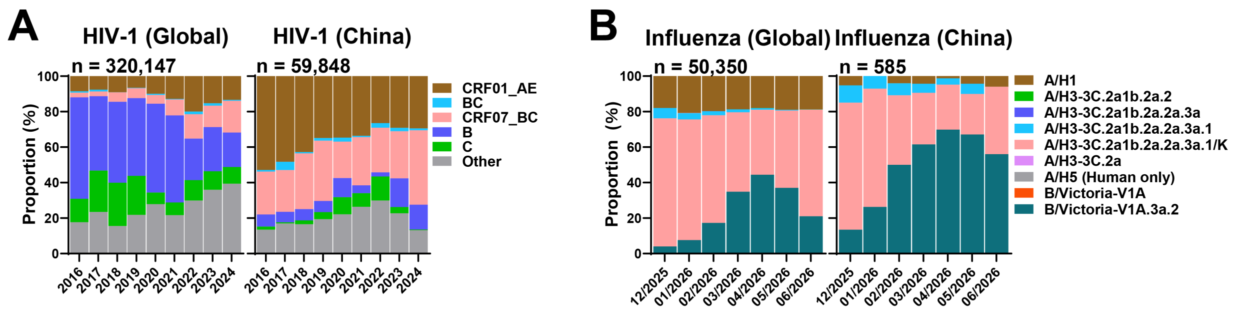


**Figure S4. Frequencies of HIV-1 and influenza virus lineages globally and in China, related to Figure 3.**

**A.** Annual proportions of major HIV-1 lineages globally and in China from 2016 to 2024. Sequences were obtained from HIV sequence databases.

**B.** Monthly proportions of major influenza virus lineages globally and in China from December 2025 to June 2026. Sequences were obtained from GISAID.

*n* indicates the total number of sequences in each dataset.

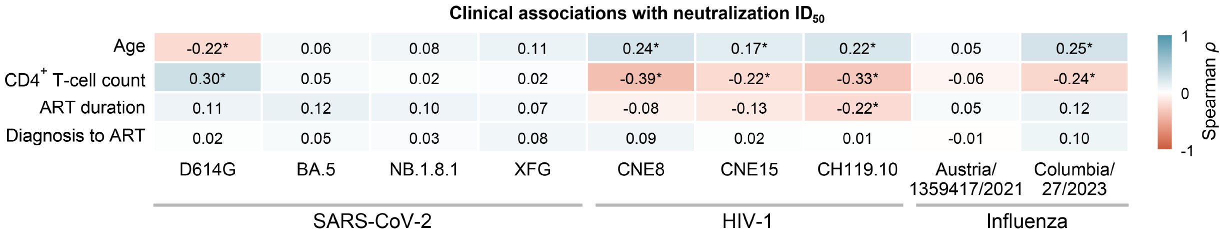


**Figure S5. Spearman correlations between clinical variables and virus-specific neutralization in PLWH, related to Figure 3.** Spearman correlations in PLWH ($n$ = 264) between age, CD4$^+$ T-cell count, ART duration, or the interval from HIV-1 diagnosis to ART initiation and $ID_{50}$ values against four SARS-CoV-2 variants, three HIV-1 pseudoviruses, and two influenza viruses. Tiles display Spearman's $\rho$, with blue and red indicating positive and negative correlations, respectively. Asterisks indicate Benjamini–Hochberg-adjusted $p < 0.05$.
